# Weaning body condition, preweaning survival, and post-weaning residence under contrasting breeding densities in Baltic grey seals

**DOI:** 10.64898/2026.09.25.754387

**Authors:** Daire Carroll, Laura Stukonytė, Karin C. Harding, Vaida Survilienė, Mart Jüssi

**Affiliations:** Department of Biological and Environmental Science, University of Gothenburg, Gothenburg, Sweden; Gothenburg Global Biodiversity Centre, Gothenburg, Sweden; Institute of Biosciences, Vilnius University, Vilnius, Lithuania; Institute of Ecology and Earth Sciences, University of Tartu, Tartu, Estonia

**Keywords:** Body condition, density dependence, life-history, population dynamics, preweaning survival, vital rates

## Abstract

In Baltic grey seals (*Halichoerus grypus*), interannual variation in sea-ice availability can alter the proportion of females breeding on land, generating substantial variation in local breeding density among years. We investigated preweaning survival, post-weaning residence, and weaning body condition at a Baltic grey seal land-breeding colony over two years with contrasting colony densities. Drone imagery provided repeated counts of pups in distinct developmental stages and estimates of moulting pup mass as an index of weaning body condition. A Bayesian state-space population model estimated substantially greater pup production in the high-(282.5 [95% CrI = 252.8 to 320.9] pups) than the low-density (188.3 [95% CrI =168.9 to 210.8] pups) year, but similar preweaning survival (0.66 [95% CrI = 0.53 to 0.81] and 0.67 [95% CrI = 0.56 to 0.79], respectively). In contrast, post-weaning residence differed substantially between the high-(11.8 [95% CrI = 9.2 to 15.7] days) and low-density (30.3 [95% CrI = 23.1 to 39.1] days) years. Greater exposure to other pups during early life was associated with lower weaning body mass (β = −0.45 [−0.89 to −0.01], P[β < 0] = 0.98). Together, these results suggest that the consequences of higher breeding density may emerge through poorer weaning condition and altered post-weaning residence even when effects on preweaning survival are not detectable.

## Introduction

Temporal variation in survival and reproductive rates determine the capacity of populations to respond to environmental change and ultimately drive fluctuations in abundance (Saether and Bakke, 2000; Morris and Doak, 2004; Rotella, 2023). Quantifying the magnitude and drivers of variation in these vital rates is central to predicting population trajectories and informing the management of wildlife populations (Saether and Bakke, 2000; Carroll et al., 2024).

Population density is a fundamental driver of variation in vital rates (Gaillard et al., 2000; Bonenfant et al., 2009). As density increases, individuals can experience greater competition for resources, increased pathogen transmission, and more frequent negative interactions with conspecifics, reducing survival and reproductive success (Fowler, 1981; Gaillard et al., 2000; Bonenfant et al., 2009).

During breeding, many pinniped species aggregate within spatially restricted colonies, creating seasonal density hotspots (Jüssi et al., 2008; Holser et al., 2021; Nagel et al., 2021). High local density can increase the occurrence of conspecific aggression, mother–pup separation, trampling, and pathogen transmission (Doidge et al., 1984; Hall et al., 2001; Stephenson et al., 2007). These processes may be particularly important during lactation, when pups are dependent on maternal care (Bowen et al., 1992; Iverson et al., 1993). For example, pup mortality from trauma and starvation increases with breeding colony density in Antarctic fur seals (*Arctocephalus gazella*) (Doidge et al., 1984). In grey seals (*Halichoerus grypus*), higher local female density has been linked to increased aggressive interactions (Stephenson et al., 2007), while pup mortality has been suggested to increase with breeding density (Coulson and Hickling, 1964; Jüssi et al., 2008).

Breeding colony density may influence offspring fitness through both lethal and sublethal pathways (Hall et al., 2001). Sublethal effects on offspring body condition may arise through reduced maternal investment leading to impaired pup development (Bowen et al., 1992; Iverson et al., 1993). Disturbance by conspecifics can reduce nursing frequency, while increased competition for prey may prolong maternal foraging trips in species that continue to forage during lactation, reducing energy transfer to offspring (Costa, 1991; Iverson et al., 1993). Consequently, pups surviving to weaning may experience impaired body condition, with potential consequences beyond the breeding season. Weaning body condition is a strong predictor of subsequent survival and lifetime fitness in many mammals (Clutton-Brock et al., 1987; Harding et al., 2005; Bowen et al., 2015). In pinnipeds, heavier pups generally have higher first-year survival and recruitment probabilities (Hall et al., 2001; Harding et al., 2005; Bowen et al., 2015).

The Baltic grey seal is a genetically isolated population with the potential to provide a valuable case study for understanding the demographic consequences of changing breeding colony density (Carroll et al., 2024; McCarthy et al., 2025). Following severe population declines caused by overhunting and pollution during the twentieth century, the population has recovered over recent decades (Harding et al., 2007; Carroll et al., 2024). Simultaneously, climate-driven reductions and northward shifts in suitable drift-ice breeding habitat are likely driving a transition from predominantly ice-breeding to land-breeding, increasing the density of breeding aggregations at terrestrial sites (Jüssi et al., 2008; Carroll et al., 2024). This shift may be further reinforced by recolonisation of the southern parts of the population’s range (Galatius et al., 2024).

Jüssi et al. (2008) compared land- and ice-breeding Baltic grey seal colonies and demonstrated that land breeding was associated with both reduced preweaning survival and poorer pup body condition. They proposed that this difference may arise from spatial constraints at terrestrial breeding sites: whereas females breeding on sea ice can spread out and maintain greater distances from neighbouring mother–pup pairs, land breeding concentrates animals within restricted areas, increasing the potential for social disturbance. However, whether variation in breeding density among terrestrial colonies is itself associated with offspring condition has remained unresolved.

Here, we tested the hypothesis that higher breeding density reduces offspring performance in Baltic grey seals. We predicted lower preweaning survival and weaning body condition at higher density. We tested these predictions over two breeding seasons with contrasting colony densities using drone-derived stage-specific counts and estimates of pup body mass.

## Methods

### Overview

Orthomosaic imagery was collected throughout the 2025 and 2026 breeding seasons at a single Baltic grey seal breeding colony. Pups were classified into developmental stages to quantify temporal changes in stage-specific abundance (Figure 1A, Appendix S1: Table S1). These observations were incorporated into a Bayesian state-space population model to estimate annual preweaning survival and post-weaning residence (Figure 1A). The same drone imagery was also used to estimate pup body mass from polygon area, providing an index of body condition at weaning (Carroll et al., 2025b). Model-derived estimates of cumulative exposure to other pups during development were then related to weaning body mass.

**Figure 1.**
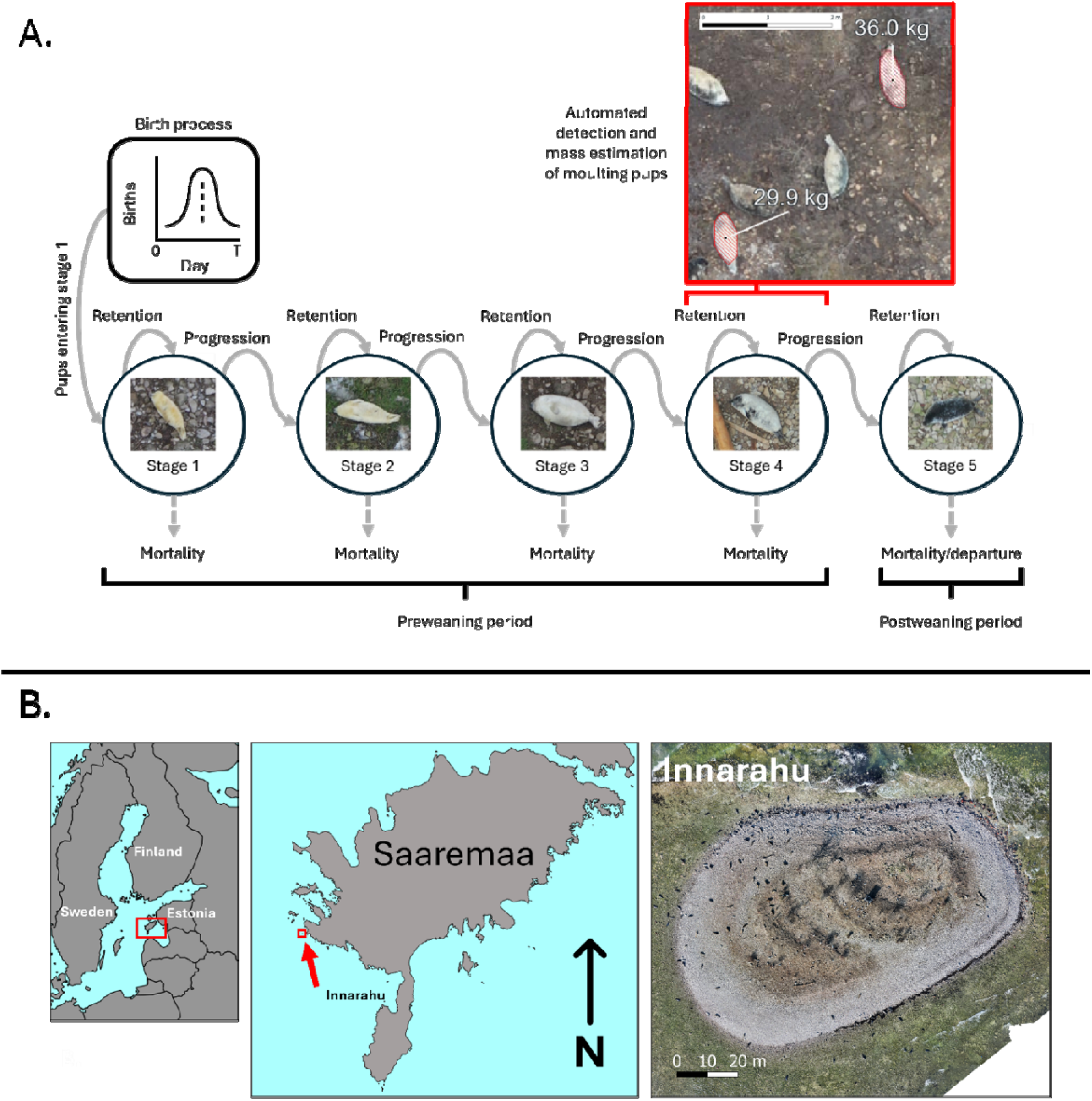
Conceptual structure of the Bayesian state-space population model for the progression of pups through developmental stages and map of the study site. **(A)** Pups enter the model through a seasonal birth process and progress through five observable developmental stages. During preweaning, individuals either remain in the same observable stage, progress to the next stage, or die. Stage 5 represents post-weaning residence. **(B)** A map indicating the study site, Innarahu, and an example of an orthomosaic. Coastline polygons sourced from Natural Earth.

### Study site

The study focused on Innarahu (58°N, 22°E), a small (approximately 8,000 m^2^) uninhabited island located less than 1 km off the west coast of Saaremaa, Estonia (Figure 1B). The island is a low-lying limestone islet consisting primarily of sand and cobble, sparse seasonal vegetation, and a small number of trees and bushes. Lunar and meteorological tides have a negligible affect on the breeding area of Innarahu due to its steep beeches and the oceanography of the Baltic Sea (Jüssi et al., 2008).

Breeding density on Innarahu varies considerably among years in response to winter sea-ice conditions (Jüssi et al., 2008). Although Innarahu itself is rarely enclosed by ice, winters with extensive sea ice in nearby sea areas provides an alternative breeding habitat, reducing the number of seals breeding on land. Conversely, during mild winters with limited sea ice, a greater proportion of the population breeds on land, substantially increasing local breeding density on Innarahu.

Surveys were carried out during the 2025 (18 February to 25 March) and 2026 (4 February to 23 March) breeding seasons. The 2025 breeding season was characterised by limited sea-ice cover in the Baltic (maximum extent = 85,000 km²), resulting in high breeding density. In contrast, the 2026 breeding season experienced substantially greater sea-ice cover (maximum extent = 181,000 km²), resulting in lower breeding colony density (Finnish Meteorological Institute, 2026).

### Data collection

Data were collected using the drone survey protocols described by Carroll et al. (2025). Weather permitting (no precipitation), daily automated drone surveys were carried out throughout the breeding season. Flights were conducted between 10:00 and 14:00 local time to maximise illumination and minimise shadowing. Images were collected using automated flight plans with 80% forward and 70% side overlap at an altitude of 45 m. Surveys were conducted using a DJI Mavic II Pro during 2025 (ground sampling distance of 1.1 cm/pixel) and a DJI Matrice 4T during 2026 (ground sampling distance of 1.2 cm/pixel). Orthomosaics were generated using WebODM (version 3.2.6), producing georeferenced imagery suitable for quantifying pup abundance, developmental stage, and body condition. Previous work has shown the comparability of different drones for orthomosaic collection (Carroll et al., 2025b). Five established developmental stages for grey seal pups (*s* = 1, 2, 3, 4, 5) (Jüssi et al., 2008; Jenssen et al., 2010) were adapted to allow classification from static orthomosaics (Figure 1B, Appendix S1: Table S1). For each survey, structured counts of pups in each developmental stage were conducted from orthomosaics using DotDotGoose (version 1.7.0) (Ersts, 2026). On the first survey of 2026 (4 February), no pups were observed. An orthomosaic was not constructed, however this date was used as the starting point during modelling.

### State space modelling

Annual preweaning survival and post-weaning residence were estimated using a novel Bayesian state-space population model fitted to the observed stage-specific pup counts (den Heyer et al., 2017; Jacobson et al., 2025) (Figure 1A, Appendix S1: Table S1). The model described the daily progression of pups through successive developmental stages using a discrete daily time step spanning the breeding season (4 February to 25 March) in each year. Hence 4 February was consistently indexed as day 1 in the model. The latent population process represented daily births, transitions among developmental stages, preweaning mortality, and post-weaning residence prior to dispersal or mortality. The observation model linked the expected number of pups in each stage to the observed survey counts. Observed stage-specific counts were modelled using a Poisson distribution, with detection probability fixed at one in the primary analysis and evaluated through subsequent sensitivity analyses. Preweaning survival and Stage 5 residence time were estimated independently for each breeding season, with identical priors applied between years. Posterior estimates were summarised as medians with 95% credible intervals (CrIs), and convergence was assessed using the potential scale reduction factor (*R□*) and effective sample size (ESS). Details of model specification, prior selection, model fitting, and sensitivity analysis are provided in Appendix S1.

### Body condition assessment

Moulting of the lanugo in grey seal pups largely coincides with weaning (Noren et al., 2008). For the purposes of this study, body condition was therefore assessed using moulting (Stage 4, Appendix S1: Table S1) pups, with estimated moulting pup mass used as an index of weaning body condition. The seal mass estimation protocol developed by Carroll et al. (2024) was adapted for grey seal pups. Moulting pups were identified from orthomosaic imagery using an automated machine-learning detector, producing spatial polygons describing the outline of each pup (Figure 1A). Polygon area was then calibrated against ground-truth body-mass measurements to estimate the mass of free-ranging moulting pups. Details of calibration and image processing are provided in Appendix S1.

The posterior median Stage 4 duration estimated by the Bayesian state-space population model was approximately one week, therefore surveys separated by this interval were selected to minimise repeated measurements of the same individuals. Three common survey dates (25 February, 5 March, and 13 March) were selected from each breeding season for analysis of moulting pup mass.

Pup density was estimated as the number of pups present per 100 m² of breeding area, assuming a constant breeding area of 6,000 m² following Jüssi et al. (2008). Pup age and date of birth were approximated from the posterior median developmental-stage durations of the state-space model, assuming that moulting pups were observed at the midpoint of Stage 4. Two cumulative density exposure metrics were then calculated from daily pup abundance, expressed as the number of pups present per 100 m² of breeding area: pup density summed over the pup’s estimated lifetime from birth to survey (cumulative lifetime density exposure), and pup density summed during early life, defined by the estimated duration of Stage 1 (cumulative early-life density exposure). Because density was summed across daily time steps, cumulative density exposure was expressed as pup-days per 100 m².

#### Analysis of trends in Stage 4 body condition

Variation in moulting pup mass was analysed using Bayesian linear models implemented in R (version 4.5.3) (R Core Team, 2024) using the *nimble* package (NIMBLE Development Team, 2026). Estimated moulting pup mass was treated as the response variable. Survey date was included as a categorical explanatory variable to account for differences in body mass among sampling periods. The two measures of cumulative density exposure were evaluated as explanatory variables in separate models: cumulative lifetime and cumulative early-life density exposure. A model containing survey date, but no density exposure term, was fitted as a null model for comparison.

Two additional sensitivity analyses were conducted for each density exposure metric. First, a survey-level random intercept was included to account for additional variation shared among pups observed during the same survey. Second, breeding season was included as a fixed effect to determine whether the estimated relationships between moulting pup mass and cumulative density exposure were robust to an overall difference in moulting pup mass between years. These sensitivity models were used to evaluate the stability of the estimated cumulative density exposure effect rather than for primary model selection.

Individual pup masses were modelled using a Gaussian error distribution. For each survey occasion, pups were assigned the posterior median cumulative density exposure estimated from the state-space population model. For each density metric, these six year-by-survey-date exposure values were standardised using their mean and standard deviation (SD) before model fitting. Normal priors were assigned to the intercept (mean = 38 kg, SD = 10 kg) and to the survey-date and cumulative density exposure regression coefficients (mean = 0, SD = 5). The residual standard deviation was assigned a uniform prior from 0 to 20 kg. For sensitivity models containing a survey-level random intercept, survey effects were assumed to arise from a normal distribution centred on zero, with their among-survey SD assigned a uniform prior from 0 to 10 kg.

Posterior distributions were sampled using three MCMC chains, each run for 100,000 iterations. The first 30,000 iterations of each chain were discarded as burn-in, no thinning was applied, and the remaining 70,000 iterations per chain were retained, yielding 210,000 posterior samples per model. Convergence was assessed using R and ESS calculated with the *coda* package (Plummer et al., 1999). Posterior estimates were summarised by the median and 95% CrI. For cumulative density exposre effects, the posterior probability of a negative relationship, *P[*β *< 0]*, was calculated. Posterior model fit was evaluated using posterior predictive checks comparing the mean and SD of observed and replicated pup masses, and Bayesian R^2^ was calculated to quantify the proportion of variation explained by the model.

The date-only, cumulative lifetime density exposure, and cumulative early-life density exposure models were compared using the widely applicable information criterion (WAIC), with WAIC weights used to describe their relative support.

## Results

### Data collection

Orthomosaics were collected on 16 survey days during the 2025 breeding season (18 February to 25 March) and 26 survey days during the 2026 breeding season (8 February to 23 March). Stage-specific pup counts from all surveys conducted in 2025 and 2026 were included in the population model. Assuming a fixed breeding area of 6000 m^2^, maximum observed pup density was higher in 2025 than in 2026, reaching 0.356 and 0.265 pups per 100 m², respectively. The earliest surveys contained only early developmental stages, with no pups observed beyond Stage 3, whereas the final surveys contained no Stage 1 or Stage 2 pups, indicating complete progression through the breeding season (Figure 2).

**Figure 2.**
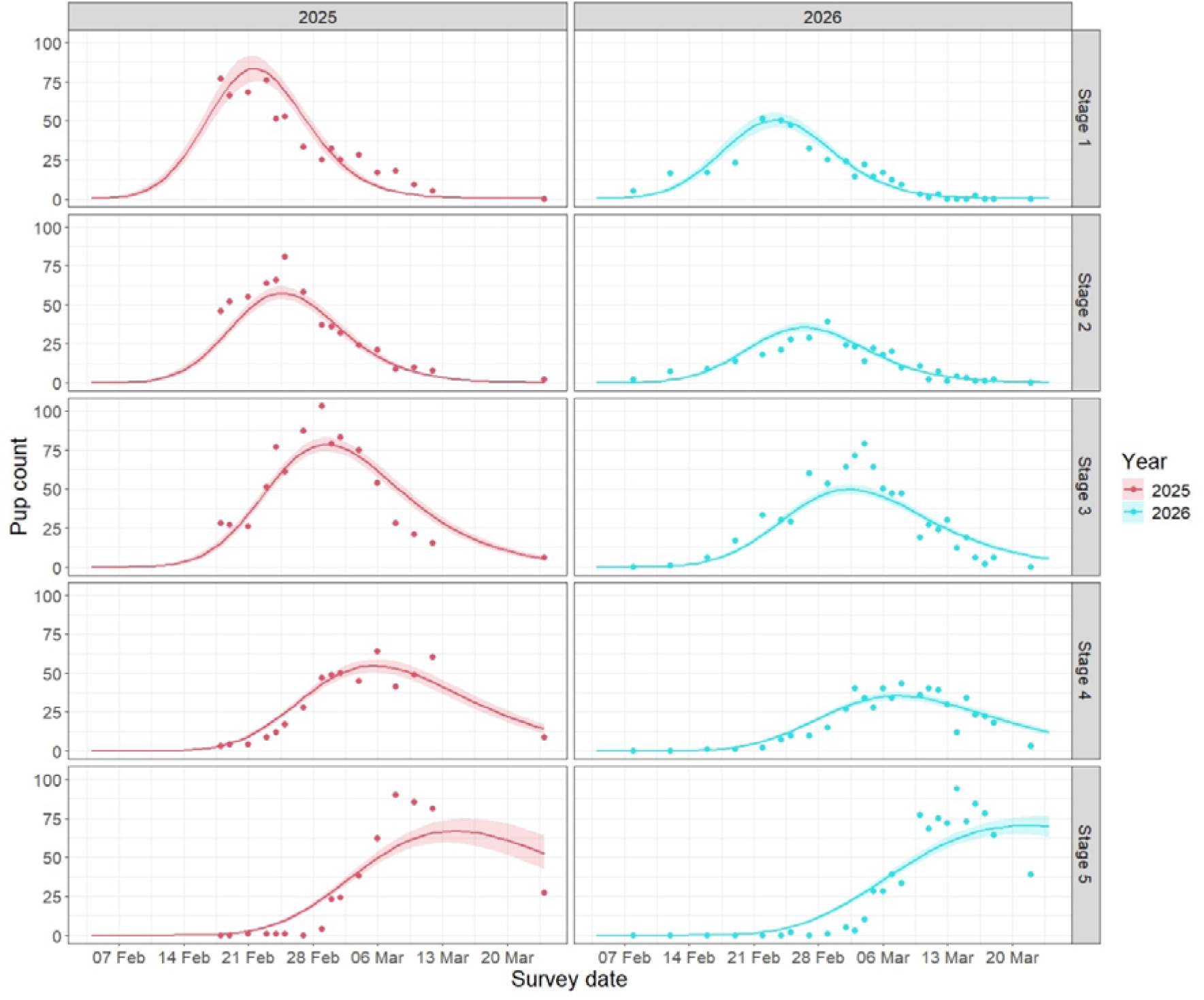
Seasonal change in number of Baltic grey seal pups in different moulting stages during two years with different ice conditions. Points represent observed stage-structured pup counts, while solid lines and shaded ribbons show the posterior median and 95% CrI of the expected number of pups in each developmental stage, as derived from a Bayesian state-space population model.

### Model performance and parameter estimates

The Bayesian state-space population model captured the overall temporal progression of developmental stages throughout both breeding seasons (Figure 2). Across all stage-specific survey counts, MAE was 7.3 pups, RMSE was 10.2 pups, and 74.8% of observations fell within the 95% posterior predictive intervals. Convergence diagnostics indicated satisfactory MCMC convergence, with all primary stochastic parameters having an R < 1.01 and a minimum ESS of 1,516. Posterior distributions (Table 1) were generally narrower than, or shifted relative to, their corresponding priors (Appendix S1: Figure S1), indicating that the observed stage-specific counts were informative for model inference.

**Table 1.** Posterior median and 95% CrIs of parameters estimated by the Bayesian state-space population model. Birth peak is expressed as model day, where day 1 corresponds to 4 February. Annual preweaning survival is reported on the probability scale.

| Parameter | Notation | Year/stage | Posterior median [95% CrI] |
| --- | --- | --- | --- |
| Total births | $B_y$ | 2025 | 282.5 [252.8 to 320.9] |
|  |  | 2026 | 188.3 [168.9 to 210.8] |
| Peak birth day | $\mu_y$ | 2025 | 16.4 [15.7 to 17.2] |
|  |  | 2026 | 18.0 [17.3 to 18.8] |
| Birth spread (days) | $\sigma_y$ | 2025 | 4.6 [4.1 to 5.2] |
|  |  | 2026 | 5.3 [4.9 to 5.6] |
| Stage duration (days) | $d_s$ | Stage 1 | 4.3 [3.9 to 4.8] |
|  |  | Stage 2 | 3.6 [3.3 to 3.8] |
|  |  | Stage 3 | 7.1 [6.7 to 7.5] |
|  |  | Stage 4 | 6.7 [6.2 to 7.3] |
| Preweaning survival | $\square_y$ | 2025 | 0.66 [0.53 to 0.81] |
|  |  | 2026 | 0.67 [0.56 to 0.79] |
| Stage 5 residence (days) | $\tau_y$ | 2025 | 11.8 [9.2 to 15.7] |
|  |  | 2026 | 30.3 [23.1 to 39.1] |

Estimated total pup production differed between years, with a posterior median difference between 2026 and 2025 of −94.6 pups (95% CrI = −126.0 to −66.6). In contrast, annual preweaning survival was similar, with a posterior median difference of 0.01 (95% CrI = −0.13 to 0.14). Post-weaning residence differed substantially between years, with a posterior median difference of 18.4 days (95% CrI = 11.3 to 26.9) (Figure 3; Table 1).

**Figure 3.**
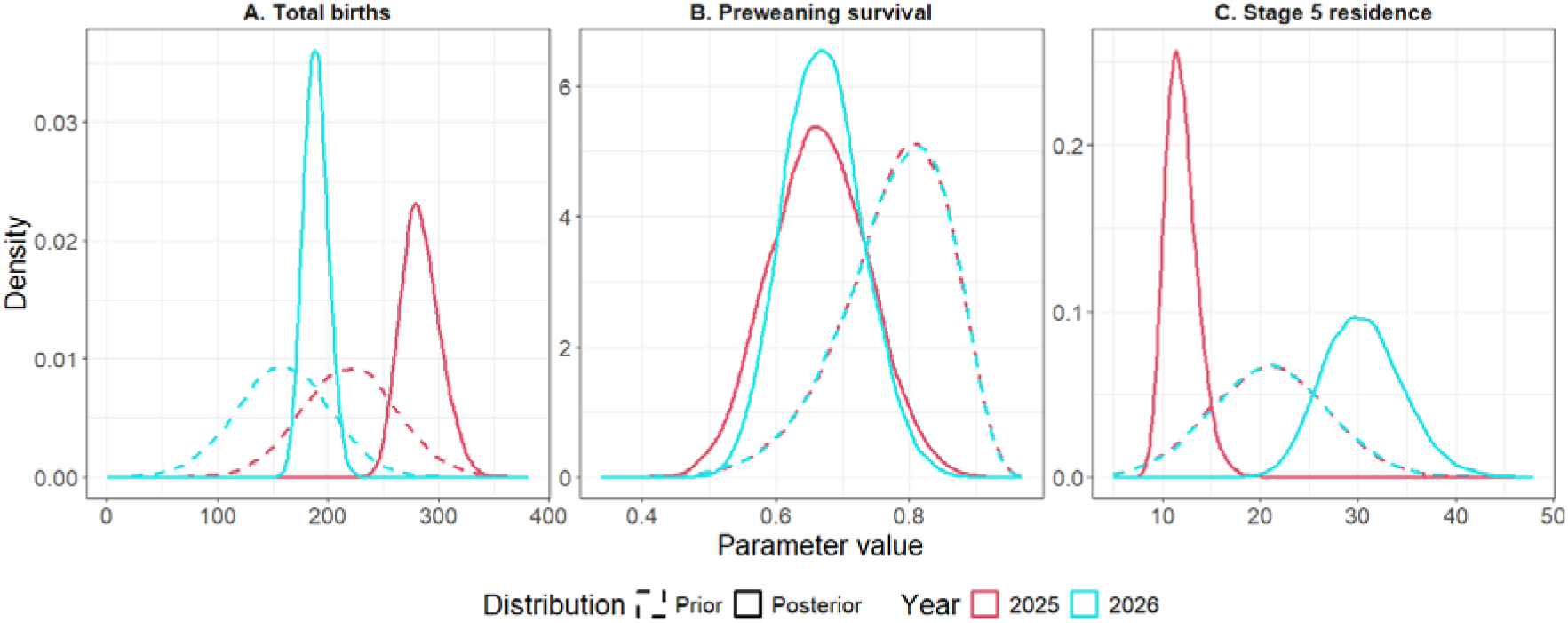
Posterior estimates of annual total births, preweaning survival, and Stage 5 residence. Prior (dashed lines) and posterior (solid lines) distributions are shown for (**A**) total births, (**B**) annual preweaning survival, and (**C**) Stage 5 residence time.

### Stage4 body condition

Across four classes (adults, white coat pups, moulting pups, and grey coat pups), the YOLOv8 segmentation model achieved a mean precision score of 0.63 and a recall score of 0.64, resulting in an F1 score of 0.63. Automated image segmentation identified 189 candidate moulting pups across the six surveys, of which 137 were retained following manual quality control (Appendix S1: Table S4). Most excluded detections reflected misclassification of other pups (42 detections), whereas misclassification of adults (3 detections), non-seal objects (1 detection) and poor polygon fits (6 detections) were comparatively uncommon. The retained dataset comprised 70 pups from 2025 and 67 pups from 2026, distributed across three survey dates in each breeding season.

Estimated moulting pup mass was modelled in relation to survey date and cumulative lifetime density exposure or cumulative early-life density exposure, with a date-only model providing the null comparison. The model including cumulative early-life density exposure received the greatest support among the three primary candidate models (WAIC = 901.67, weight = 0.55). The cumulative lifetime density exposure model received somewhat less support (WAIC = 902.87, weight = 0.30), while the date-only null model received the least support (WAIC = 904.36, weight = 0.14). Differences in model support were modest, but overall favoured the inclusion of cumulative density exposure, with greater support for early-life than lifetime exposure.

In the cumulative early-life density exposure model, estimated Stage 4 pup mass declined with increasing density exposure (β = −0.45 [95% CrI −0.89 to −0.01]; P[β < 0] = 0.98; Figure 4A and B). The estimated relationship was similar in magnitude when a survey-level random intercept was included (β = −0.40 [95% CrI = −1.03 to 0.29]; P[β < 0] = 0.90) and when breeding season was included as a fixed effect (β = −0.43 [95% CrI = −1.03 to 0.18; P[β < 0] = 0.92), although posterior uncertainty increased in both sensitivity analyses.

**Figure 4.**
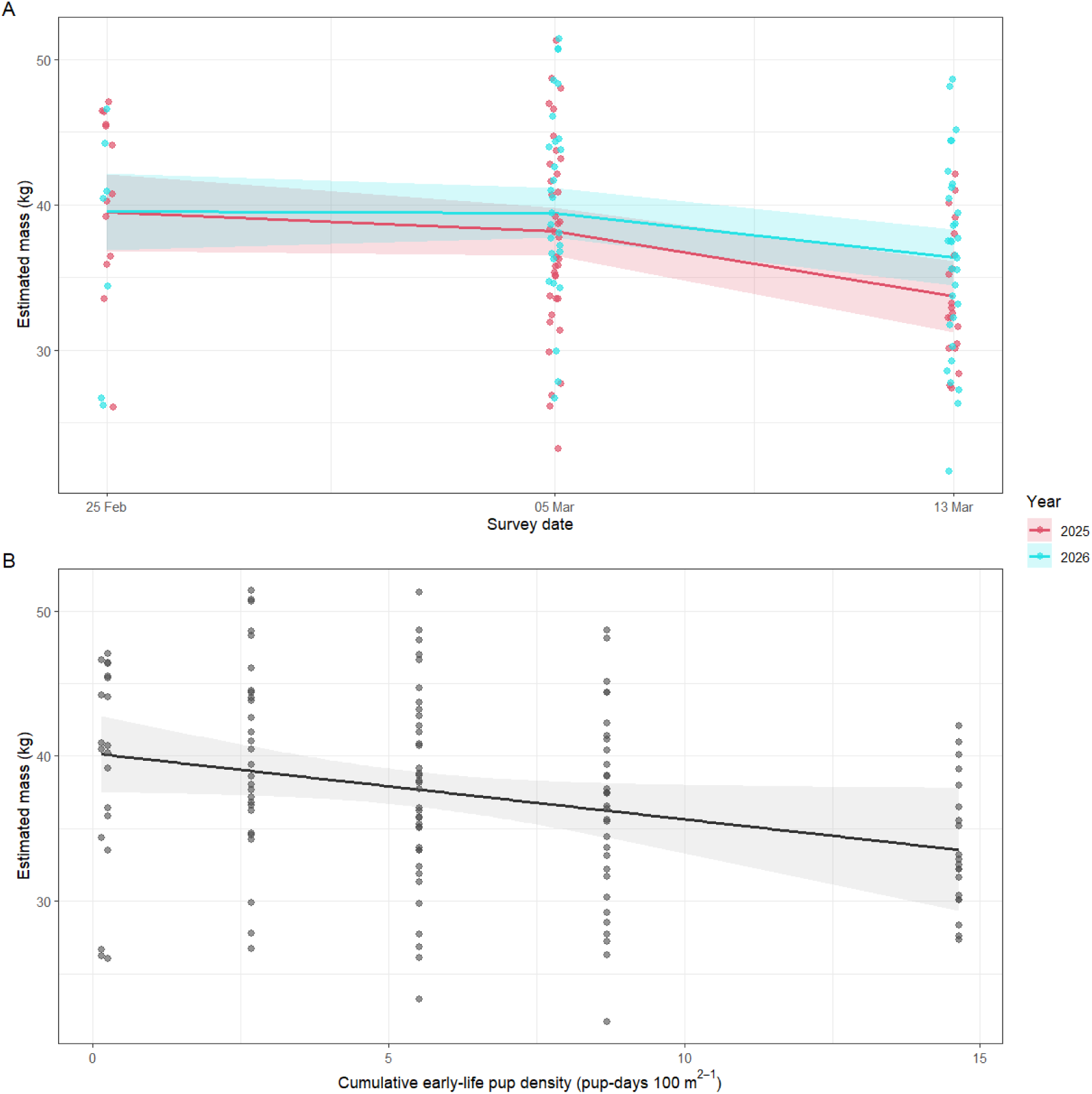
Relationships between moulting pup mass, survey date and cumulative density exposure. (**A**) Estimated moulting pup mass across the shared survey period in 2025 and 2026. Solid lines show posterior median predictions from the cumulative early-life density exposure model and shaded ribbons show 95% CrIs; points represent individual estimated pup masses. (**B**) Predicted relationship between moulting pup mass and cumulative early-life density exposure, expressed as pup-days per 100 m². The solid line shows the posterior median model prediction, and the shaded ribbon shows the 95% CrI. Points represent individual estimated pup masses plotted against the posterior median cumulative early-life density exposure assigned to their survey occasion.

The primary model explained a relatively small proportion of variation in individual pup mass (Bayesian R^2^ = 0.10 [95% CrI = 0.03 to 0.20]), however, posterior predictive distributions closely reproduced the observed mean mass (observed = 37.5 kg; posterior predictive median = 37.5 kg [95% interval = 36.0 to 39.1]) and standard deviation (observed = 6.63 kg; posterior predictive median = 6.75 kg [95% interval = 5.71 to 8.00]). MCMC diagnostics indicated satisfactory convergence (maximum R < 1.01; minimum ESS = 10,821).

Predicted mean mass on 25 February was similar between breeding seasons, at approximately 39.5 kg in both 2025 and 2026. By 13 March, predicted mean mass had declined to 33.7 kg in 2025 and 36.4 kg in 2026 (Figure 4A). This represented predicted declines of 5.8 kg (95% CrI = 2.0 to 9.6 kg) and 3.2 kg (95% CrI = 0.01 to 6.2 kg), respectively, across the shared sampling period (Figure 4B).

## Discussion

### Summary

We investigated interannual variation in lethal and sublethal components of offspring fitness in Baltic grey seals during two breeding seasons with contrasting colony densities. The high-density season was associated with poorer body condition at weaning, while preweaning survival was similar between years. Across both breeding seasons, weaning body mass was negatively associated with cumulative density exposure experienced during early life, providing evidence consistent with a sublethal density-dependent response. Post-weaning residence also differed substantially between breeding seasons, indicating additional interannual variation in pup behaviour or mortality after weaning.

### Evaluation of methods

Systematic aerial surveys have formed the basis of pinniped population monitoring for several decades. Traditional approaches have relied on repeated aerial photography followed by manual interpretation of imagery (Teilmann et al., 2010; Carroll et al., 2024, 2025a). More recently, advances in drone technology have substantially improved the spatial resolution, flexibility, and accessibility of aerial surveys (Infantes et al., 2022; Amorosi et al., 2024; Carroll et al., 2025b). By combining repeated drone surveys with Bayesian state-space modelling and automated image analysis, we simultaneously estimated annual preweaning survival, post-weaning residence, and body condition within a breeding colony. These complementary measures provide substantially greater ecological information than pup counts alone.

State-space models have increasingly been adopted to estimate various aspects of pinniped population dynamics from aerial survey data, providing a flexible framework for combining biological processes with repeated observations. Previous models have been developed to estimate pup production, birth phenology, or population size (Lonergan et al., 2011; Thomas et al., 2019; Mosnier et al., 2023; Jacobson et al., 2025). Our model addresses a different ecological question by incorporating observed developmental stages and an explicit survival process within the latent population model. Possible extensions of our model are to investigate stage specific survival, or variation in survival across the breeding season.

Our stage structured counts are subject to several sources of uncertainty. Although aerial imagery provided good visibility of the island, some pups were likely missed when partially obscured by vegetation, resulting in incomplete detection. Developmental stage classification inevitably contains some subjectivity because stages represent a continuous biological process rather than discrete categories (Jüssi et al., 2008; Jenssen et al., 2010). Individual variation in development may violate the assumption that all pups pass sequentially through the defined stage durations. For example, pups in poor body condition may begin moulting before taking on the characteristic Stage 3 body shape or otherwise retaining the size of a Stage 1 of 2 pup, lengthening the apparent duration of these stages. The model accounts only for pups that enter the observable population process. Stillborn pups or pups dying shortly after birth may be missed between surveys, particularly because small carcasses can be rapidly scavenged or removed by large birds, including eagles and gulls that occur in high numbers at the site. Consequently, early pup mortality may be underestimated due to imperfect detection. Post-weaning residence represents the time Stage 5 pups remain observable at the colony; because individuals were not tracked in the current study, exit following weaning cannot be separated into dispersal and post-weaning mortality. Consequently, estimated preweaning survival should be interpreted as survival within the observed stage-structured population rather than a complete measure of all perinatal mortality.

The two-year study design limits attribution of interannual differences specifically to breeding density. Pup abundance differed markedly between years, but other environmental conditions also differed and could have influenced pup development or maternal investment. For example, ice conditions, temperature, or precipitation could contribute to differences in pup body condition or post-weaning behaviour. The observed negative relationship between cumulative density exposure and pup mass therefore provides evidence consistent with a density-dependent response, but the present study cannot fully separate density from other year-specific environmental effects. Replication across additional breeding seasons spanning a wider range of colony densities and environmental conditions would provide a stronger test of causality.

Our metrics of breeding colony density were based on pup abundance only. It is likely that the number of adults, including males, is an important factor driving changes in pup survival and body condition (Hall et al., 2001; Langley et al., 2026). Although adults could be observed in our images and male and female adults could largely be distinguished, they were more mobile than pups and spent a large amount of time in the water and so counts were less stable than for pups which only rarely enter the water before moulting (Jüssi et al., 2008; Jenssen et al., 2010). Pup abundance should therefore be interpreted as a consistent index of colony density rather than a complete measure of the social environment experienced by pups.

Despite these limitations, several lines of evidence support the reliability of the estimated demographic patterns. The state-space model closely reproduced the observed progression of developmental stages throughout both breeding seasons, convergence diagnostics indicated stable parameter estimation, and posterior estimates were generally robust to alternative prior specifications. Variation in assumed detection probability modestly affected the absolute estimates of preweaning survival and post-weaning residence but did not alter the interannual patterns. Across sensitivity analyses, preweaning survival remained similar between years, whereas post-weaning residence was consistently longer in 2026. Consequently, although the absolute magnitude of these parameters should be interpreted with some caution, the contrasting patterns in survival and post-weaning residence were consistently supported by the observed stage-structured counts.

Machine-learning approaches are becoming an increasingly important component of wildlife monitoring, substantially reducing the time required to process large image datasets (Pichler and Hartig, 2023; Carroll et al., 2025b). Previous work has demonstrated that automated detection can achieve high accuracy for pinniped surveys, allowing drone imagery to be processed efficiently for estimates of abundance, reproduction, and body size (Infantes et al., 2022; Carroll et al., 2025b). The present study demonstrates that satisfactory performance can be achieved using an entirely open-source workflow. Although the segmentation model occasionally misclassified moulting pups as adjacent developmental stages, confusion between adults and pups or false detections of non-seal objects were rare. Importantly, all automated detections were subjected to manual quality control before analysis, ensuring that body-mass estimates were based only on correctly identified Stage 4 pups. Such manual controls may not be required if coarser census counts or fewer classifications are sufficient (Thomas et al., 2019; Jacobson et al., 2025).

Our body-mass estimates compare favourably with previously published photogrammetric approaches (Shero et al., 2021; Carroll et al., 2025b). The simple relationship between polygon area and measured body mass produced a mean absolute prediction error of 3.4 kg. This level of accuracy is comparable to the volumetric approach developed by Carroll et al. (2025), which reported mean absolute errors of approximately 3.2 kg for harbour seal pups, despite requiring substantially more complex measurements of body length, width and estimated volume.

### Ecological relevance

As capital breeders investing substantial stored energy in a single pup, grey seal reproductive fitness depends not only on offspring production but also on offspring quality (McNamara and Houston, 1996; Jüssi et al., 2008). Body condition at weaning is widely recognised as a strong predictor of fitness in seals (Hall et al., 2001; Harding et al., 2005; Bowen et al., 2015). Despite the recognised importance of weaning condition, relatively little evidence has linked these fitness consequences directly to breeding colony density. Previous studies have shown that increasing local density can elevate aggressive interactions among breeding females and increase the risk of mother–pup separation, transmission of various pathogens, trampling, and starvation, yet direct relationships between colony density and pup mortality have proved difficult to demonstrate (Coulson and Hickling, 1964; Hall et al., 2001; Twiss et al., 2003; Ashley et al., 2020). The model containing cumulative exposure to other pups during early life received greater support than the model representing cumulative density over the pup’s estimated lifetime, although the difference in support was modest. This is consistent with the hypothesis that the early lactation period is a key window during which density-dependent processes influence offspring fitness (Iverson et al., 1993). Although the underlying mechanism cannot be identified from the present study, increased social disturbance, competition for nursing space, and mother–pup separation all represent plausible pathways linking elevated breeding colony density to reduced pup growth. These mechanisms could be explored through future behavioural studies.

The substantial interannual difference in post-weaning residence provides a further potential link between breeding conditions and offspring performance. Stage 5 residence was considerably shorter during the higher-density 2025 breeding season than in 2026. At weaning, approximately 40–50% of grey seal pup body mass comprises fat, which provides the principal energy reserve supporting the post-weaning fast (Reilly, 1991; Noren et al., 2008). Body mass and composition at weaning can therefore influence the duration of this period and the timing of departure to sea (Muelbert et al., 2003; Noren et al., 2008). Although we did not directly measure individual metabolic transitions or departure, the shorter residence of fully moulted pups during the higher-density season is consistent with this energetic framework. A plausible explanation for the shorter Stage 5 residence observed in 2025 is that poorer weaning condition resulted in more rapid depletion of energy reserves and earlier departure from the colony. This interpretation is consistent with our finding of lower body mass with increasing cumulative density exposure. The duration of the post-weaning period may have important consequences for subsequent performance, because this period contributes to the development of physiological capacities required for independent foraging, including muscle development, myoglobin stores and diving capacity (Bennett et al., 2007, 2010; Noren et al., 2008). Consequently, an association between breeding colony density, weaning condition, and post-weaning residence provides a plausible pathway through which sublethal effects experienced during lactation could extend beyond the breeding period.

Our findings are consistent with those of Jüssi et al. (2008), who demonstrated that breeding habitat and local breeding density strongly influenced pup body condition in Baltic grey seals. They reported a mean preweaning mortality of 21.1% for land-breeding colonies, compared with only 1.5% for pups born on sea ice (based on recorded number of dead pups, without a sound estimate of total births). They further found that pups reared on ice were substantially heavier at the onset of moult (48.3 ± 8.1 kg) than those reared on land (37.4 ± 7.8 kg). They proposed that the higher breeding densities associated with land breeding reduce offspring performance through increased social disturbance. Our mean estimated moulting pup masses of 37.1 kg in 2025 and 38.0 kg in 2026 are similar to the mean mass reported by Jüssi et al. (2008) for land-breeding pups. Within this range of land-breeding conditions, we found that weaning mass declined with increasing cumulative colony density exposure during early life. Importantly, this relationship was estimated across repeated sampling occasions within the two breeding seasons rather than from the annual contrast alone, providing evidence consistent with the proposed sublethal density-dependent response without the potentially confounding comparison between ice- and land-breeding habitats.

In contrast, estimated preweaning survival was similar between breeding seasons despite substantially higher pup production and breeding density in 2025. Our estimates of approximately 0.66 to 0.67 were substantially lower than the mean preweaning survival of 0.79 reported by Jüssi et al. (2008) for Baltic land-breeding colonies, although their mortality estimates were based on recorded dead pups and are therefore not directly comparable with the stage-structured survival estimates used here. The relatively low survival estimated at Innarahu should not be interpreted as representative of land-breeding Baltic grey seals more generally (Carroll et al., 2024; Vanko et al., 2026). Innarahu supports a particularly dense breeding aggregation, whereas terrestrial breeding sites elsewhere differ substantially in colony size and density. Our results instead suggest that variation in breeding density may be detectable in offspring condition even when a corresponding difference in preweaning survival is not evident. Such sublethal responses may provide a more sensitive indicator of variation in breeding conditions than mortality alone.

### Conclusion

Higher breeding colony density was associated with poorer body condition at weaning in Baltic grey seals, despite similar preweaning survival between years. By combining repeated drone surveys with Bayesian state-space modelling and automated body-mass estimation, we quantified demographic and phenotypic components of offspring performance from the same aerial imagery. The negative relationship between cumulative early-life colony density and weaning mass provides evidence consistent with a sublethal density-dependent response, suggesting that variation in breeding conditions may be expressed through offspring condition without a corresponding detectable difference in mortality. Integrating measures of preweaning survival and weaning condition into long-term monitoring could therefore provide a more complete assessment of population productivity and its response to changing breeding conditions.

## Data accessibility statement

All data and code are available at Zenodo.org (DOI: 10.5281/zenodo.22748080).

## Statement of authorship

D. Carroll contributed to study conception, developed the methodology, collected the data, conducted the analyses, and led the writing of the manuscript. M. Jüssi contributed to study conception and data collection. Laura Stukonytė and Vaida Survilienė contributed to data collection. Karin C. Harding contributed to study conception. All authors contributed to the interpretation of the results, reviewed and revised the manuscript, and approved the final version.

## Acknowledgments

The authors thank I. Kučinskaitė-Kodzė, M. Babonaitė, T. Vainauskas, K. Lenko and numerous students from the Vilnius University Grey Seal Research Group for their invaluable help in capturing, weighing, and marking animals in the field. The authors thank J. Harvey-Carroll for assistance with data collection and her comments on the manuscript. D. Carroll acknowledges grants from the Wild Animal Initiative (F-2023-00005), and the Swedish Environmental Protection Agency through the Environmental Research Fund and the Swedish Research Council Formas (2024-00147).

## Permits

Flights and animal handling were conducted under permission from the Estonian Environmental Board (permit nr. 1/3/20/60).

## References

Amorosi, L., Carroll, D., Carroll, P. & Esposito Amideo, A. (2024). Optimal drone routing for seal pup counts. In: Optimization in Green Sustainability and Ecological Transition (eds. Bruglieri, M., Festa, P., Macrina, G. & Pisacane, O.). Springer Nature Switzerland, Cham, pp. 147–156.

Ashley, E.A., Olson, J.K., Adler, T.E., Raverty, S., Anderson, E.M., Jeffries, S. et al. (2020). Causes of mortality in a harbor seal (*Phoca vitulina*) population at equilibrium. Front. Mar. Sci., 7, 319.

Bennett, K.A., McConnell, B.J., Moss, S.E.W., Speakman, J.R., Pomeroy, P.P. & Fedak, M.A. (2010). Effects of age and body mass on development of diving capabilities of gray seal pups: costs and benefits of the postweaning fast. Physiol. Biochem. Zool., 83, 911–923.

Bennett, K.A., Speakman, J.R., Moss, S.E.W., Pomeroy, P. & Fedak, M.A. (2007). Effects of mass and body composition on fasting fuel utilisation in grey seal pups (*Halichoerus grypus* Fabricius): an experimental study using supplementary feeding. J. Exp. Biol., 210, 3043– 3053.

Bonenfant, C., Gaillard, J.-M., Coulson, T., Festa-Bianchet, M., Loison, A., Garel, M. et al. (2009). Empirical evidence of density-dependence in populations of large herbivores. In: *Advances in Ecological Research*, 41, pp. 313–357.

Bowen, W.D., Stobo, W.T. & Smith, S.J. (1992). Mass changes of grey seal *Halichoerus grypus* pups on Sable Island: differential maternal investment reconsidered. J. Zool., 227, 607–622.

Bowen, W.D., den Heyer, C.E., McMillan, J.I. & Iverson, S.J. (2015). Offspring size at weaning affects survival to recruitment and reproductive performance of primiparous gray seals. Ecol. Evol., 5, 1412–1424.

Carroll, D., Ahola, M.P., Carlsson, A.M., Galatius, A., Nilssen, K.T., Härkönen, T. et al. (2025a). Declining harbour seal abundance in a previously recovering meta-population. PLoS ONE, 20, e0326933.

Carroll, D., Ahola, M.P., Carlsson, A.M., Sköld, M. & Harding, K.C. (2024). 120-years of ecological monitoring data shows that the risk of overhunting is increased by environmental degradation for an isolated marine mammal population: the Baltic grey seal. J. Anim. Ecol., 93, 525–539.

Carroll, D., Infantes, E., Pagan, E.V. & Harding, K.C. (2025b). Approaching a population-level assessment of body size in pinnipeds using drones, an early warning of environmental degradation. *Remote Sens*. Ecol. Conserv., 11, 156–171.

Clutton-Brock, T.H., Major, M., Albon, S.D. & Guinness, F.E. (1987). Early development and population dynamics in red deer. I. Density-dependent effects on juvenile survival. J. Anim. Ecol., 56, 53–67.

Costa, D.P. (1991). Reproductive and foraging energetics of pinnipeds: implications for life history patterns. In: The Behaviour of Pinnipeds (ed. Renouf, D.). Springer Netherlands, Dordrecht, pp. 300–344.

Coulson, J.C. & Hickling, G. (1964). The breeding biology of the grey seal, *Halichoerus grypus* (Fab.), on the Farne Islands, Northumberland. J. Anim. Ecol., 33, 485–512.

Doidge, D.W., Croxall, J.P. & Baker, J.R. (1984). Density-dependent pup mortality in the Antarctic fur seal *Arctocephalus gazella* at South Georgia. J. Zool., 202, 449–460.

Ersts, P.J. (2026). DotDotGoose. Available at: https://biodiversityinformatics.amnh.org/open_source/dotdotgoose/. Last accessed 31 JULY 2026.

Finnish Meteorological Institute. (2026). Ice season 2025–2026. Available at: https://en.ilmatieteenlaitos.fi/ice-winter-2025-2026. Last accessed 29 JULY 2026.

Fowler, C.W. (1981). Density dependence as related to life history strategy. Ecology, 62, 602–610.

Gaillard, J.-M., Festa-Bianchet, M., Yoccoz, N.G., Loison, A. & Toïgo, C. (2000). Temporal variation in fitness components and population dynamics of large herbivores. Annu. Rev. Ecol. Syst., 31, 367–393.

Galatius, A., Olsen, M.T., Allentoft-Larsen, M., Balle, J.D., Kyhn, L.A., Sveegaard, S. et al. (2024). Evidence of distribution overlap between Atlantic and Baltic grey seals. J. Mar. Biol. Assoc. U.K., 104, e30.

Hall, A.J., McConnell, B.J. & Barker, R.J. (2001). Factors affecting first-year survival in grey seals and their implications for life history strategy. J. Anim. Ecol., 70, 138–149.

Harding, K.C., Fujiwara, M., Axberg, Y. & Härkönen, T. (2005). Mass-dependent energetics and survival in harbour seal pups. Funct. Ecol., 19, 129–135.

Harding, K.C., Härkönen, T., Helander, B. & Karlsson, O. (2007). Status of Baltic grey seals: population assessment and extinction risk. NAMMCO Sci. Publ., 6, 33–56.

Holser, R.R., Crocker, D.E., Robinson, P.W., Condit, R. & Costa, D.P. (2021). Density-dependent effects on reproductive output in a capital breeding carnivore, the northern elephant seal (*Mirounga angustirostris*). Proc. R. Soc. B, 288, 20211258.

Infantes, E., Carroll, D., Silva, W.T.A.F., Härkönen, T., Edwards, S.V. & Harding, K.C. (2022). An automated work-flow for pinniped surveys: a new tool for monitoring population dynamics. Front. Ecol. Evol., 10, 905309.

Iverson, S.J., Bowen, W.D., Boness, D.J. & Oftedal, O.T. (1993). The effect of maternal size and milk energy output on pup growth in grey seals (*Halichoerus grypus*). Physiol. Zool., 66, 61–88.

Jacobson, E.K., Goldman, M.R., Thomas, L. & Russell, D.J.F. (2025). A state-space model for estimating pinniped pup production from serial counts at breeding colonies. Ecol. Modell., 510, 111333.

Jenssen, B.M., Åsmul, J.I., Ekker, M. & Vongraven, D. (2010). To go for a swim or not? Consequences of neonatal aquatic dispersal behaviour for growth in grey seal pups. Anim. Behav., 80, 667–673.

Jüssi, M., Härkönen, T., Helle, E. & Jüssi, I. (2008). Decreasing ice coverage will reduce the breeding success of Baltic grey seal (*Halichoerus grypus*) females. Ambio, 37, 80–85.

Langley, I., Lidgard, D., Varkey, P., Sanchez, M., Rivard, M. & den Heyer, C.E. (2026). Gray seal cannibalism at the largest colony in the world, Sable Island. Mar. Mamm. Sci., 42, e70138.

Lonergan, M., Thompson, D., Thomas, L. & Duck, C. (2011). An approximate Bayesian method applied to estimating the trajectories of four British grey seal (*Halichoerus grypus*) populations from pup counts. J. Mar. Biol., 2011, 1–7.

McCarthy, M.L., Cammen, K.M., Granquist, S.M., Dietz, R., Teilmann, J., Thøstesen, C.B. et al. (2025). Range-wide genomic analysis reveals regional and meta-population dynamics of decline and recovery in the grey seal. Mol. Ecol., 34, e17824.

McNamara, J.M. & Houston, A.I. (1996). State-dependent life histories. Nature, 380, 215– 221.

Morris, W.F. & Doak, D.F. (2004). Buffering of life histories against environmental stochasticity: accounting for a spurious correlation between the variabilities of vital rates and their contributions to fitness. Am. Nat., 163, 579–590.

Mosnier, A., den Heyer, C.E., Stenson, G.B. & Hammill, M.O. (2023). A Bayesian birth distribution model for grey seals and an evaluation of the timing of harvest. DFO Can. Sci. Advis. Sec.

Muelbert, M.M.C., Bowen, W.D. & Iverson, S.J. (2003). Weaning mass affects changes in body composition and food intake in harbour seal pups during the first month of independence. Physiol. Biochem. Zool., 76, 418–427.

Nagel, R., Stainfield, C., Fox-Clarke, C., Toscani, C., Forcada, J. & Hoffman, J.I. (2021). Evidence for an Allee effect in a declining fur seal population. Proc. R. Soc. B, 288, 20202882.

NIMBLE Development Team. (2026). NIMBLE: MCMC, particle filtering, and programmable hierarchical modeling. Zenodo. Available at: 10.5281/zenodo.1211190.

Noren, S.R., Boness, D.J., Iverson, S.J., McMillan, J. & Bowen, W.D. (2008). Body condition at weaning affects the duration of the postweaning fast in gray seal pups (*Halichoerus grypus*). Physiol. Biochem. Zool., 81, 269–277.

Pichler, M. & Hartig, F. (2023). Machine learning and deep learning—a review for ecologists. Methods Ecol. Evol., 14, 994–1016.

Plummer, M., Best, N., Cowles, K., Vines, K., Sarkar, D., Bates, D. et al. (1999). coda: output analysis and diagnostics for MCMC. R package.

R Core Team. (2024). R: A language and environment for statistical computing. R Foundation for Statistical Computing, Vienna. Available at: https://www.R-project.org/.

Reilly, J.J. (1991). Adaptations to prolonged fasting in free-living weaned gray seal pups. Am. J. Physiol. Regul. Integr. Comp. Physiol., 260, R267–R272.

Rotella, J.J. (2023). Patterns, sources, and consequences of variation in age-specific vital rates: insights from a long-term study of Weddell seals. J. Anim. Ecol., 92, 552–567.

Sæther, B.-E. & Bakke, O. (2000). Avian life history variation and contribution of demographic traits to the population growth rate. Ecology, 81, 642–653.

Shero, M.R., Dale, J., Seymour, A.C., Hammill, M.O., Mosnier, A., Mongrain, S. et al. (2021). Tracking wildlife energy dynamics with unoccupied aircraft systems and three-dimensional photogrammetry. Methods Ecol. Evol., 12, 2458–2472.

Stephenson, C.M., Matthiopoulos, J. & Harwood, J. (2007). Influence of the physical environment and conspecific aggression on the spatial arrangement of breeding grey seals. Ecol. Inform., 2, 308–317.

Teilmann, J., Rigét, F. & Härkönen, T. (2010). Optimizing survey design for Scandinavian harbour seals: population trend as an ecological quality element. ICES J. Mar. Sci., 67, 952– 958.

Thomas, L., Russell, D.J.F., Duck, C.D., Morris, C.D., Lonergan, M., Empacher, F. et al. (2019). Modelling the population size and dynamics of the British grey seal. Aquat. Conserv., 29, 6–23.

Twiss, S.D., Duck, C. & Pomeroy, P.P. (2003). Grey seal (*Halichoerus grypus*) pup mortality not explained by local breeding density on North Rona, Scotland. J. Zool., 259, 83–91.

Vanko, M., Helle, I., Kunnasranta, M., Ahola, M.P., Bäcklin, B.-M., Carlsson, A.M. et al. (2026). Bayesian integrated population model of Baltic grey seals (*Halichoerus grypus grypus*) informs on population carrying capacity and the impact of Baltic herring (*Clupea harengus membras*) on reproduction. Ecol. Modell., 519, 111666.

