## Appendix S1 for "Weaning body condition, preweaning survival, and post-weaning residence under contrasting breeding densities in Baltic grey seals"

Appendix S1: Weaning body condition, preweaning survival, and post-weaning residence under contrasting breeding densities in Baltic grey seals

Daire Carroll^1,2,*^, Laura Stukonytė^3^, Karin C. Harding^1^, Vaida Survilienė^3^, Mart Jüssi^4^

^1^ Department of Biological and Environmental Science, University of Gothenburg, Gothenburg, Sweden

^2^ Gothenburg Global Biodiversity Centre, Gothenburg, Sweden

^3^ Institute of Biosciences, Vilnius University, Vilnius, Lithuania

^4^ Institute of Ecology and Earth Sciences, University of Tartu, Tartu, Estonia

### Bayesian state-space population modelling

#### Model specification

The breeding season was represented using a discrete daily time step, indexed by *t = 1, ..., T*, where $T$ corresponded to the period from 4 February to 25 March. The 2025 and 2026 breeding seasons were modelled jointly, with year indexed by *y*. The latent state variable, *N_y,t,s_*, represented the expected number of pups in developmental stage *s* on day *t* of breeding season $y$. Model dynamics were described by four biological processes: birth, developmental-stage progression, preweaning survival, and post-weaning residence.

Daily births were modelled using a Gaussian-shaped birth curve. The annual total number of births, $B_{y}$, was distributed through time according to a Gaussian distribution centred on the peak birth day, $\mu_{y}$, with birth-season spread, $\sigma_{y}$,

$$w_{y,t}=\exp\left[ -\frac{1}{2}\left( \frac{t-\mu_{y}}{\sigma_{y}} \right)^{2} \right],$$

Where *w_y,t_* is the unnormalised birth weight for day *t*. Expected daily births ($b_{y,t}$) were obtained by normalising these weights,

$$b_{y,t}=B_{y}\frac{w_{y,t}}{\sum_{t=1}^{T} w_{y,t}},$$

such that daily births summed to the annual total,

$$\sum_{t=1}^{T} b_{y,t}=B_{y}.$$

Progression through developmental stages was represented as a geometric residence process. The mean stage duration, *d_s_*, denoted the mean duration in days of developmental stage $s$. The corresponding daily progression probability, *γ*_s_, was

$$\gamma_{s}=\frac{1}{d_{s}}.$$

The annual preweaning survival probability, *ϕ_y_*, represented survival from birth until completion of Stage 4 and was estimated independently for each breeding season. Survival was partitioned among developmental stages in proportion to their estimated durations. If the total preweaning duration, *D*, was

$$D=\sum_{s=1}^{4} d_{s},$$

then the stage-duration proportion, *ω_s_* represented the proportion of the total preweaning period spent in developmental stage $s$,

$$\omega_{s}=\frac{d_{s}}{D},$$

giving the stage-completion survival probability

$$\phi_{y,s}^{*}=\left( \phi_{y} \right)^{\omega_{s}}.$$

The daily survival probability *q_y,s_* was then derived from the geometric residence process as

$$q_{y,s}=\frac{\phi_{y,s}^{*}}{\phi_{y,s}^{*}(1-\gamma_{s})+\gamma_{s}},$$

which yielded the daily probability of remaining within a stage,

$$r_{y,s}=q_{y,s}(1-\gamma_{s}),$$

or the daily probability of progressing to the next stage,

$$a_{y,s}=q_{y,s}\gamma_{s}.$$

The daily mortality probability, *m_y,s_* within each developmental stage was therefore

$$m_{y,s}=1-r_{y,s}-a_{y,s}.$$

Stage 5 represented post-weaning residence in the colony rather than an additional developmental stage. The annual mean Stage 5 residence time (*τ_y_*) denoted the average number of days that recently weaned pups remained ashore before departing in breeding season *y*. Stage 5 residence was estimated independently for each breeding season. The corresponding annual daily retention probability, *ρ_y_*, was

$$\rho_{y}=1-\frac{1}{\tau_{y}},$$

such that pups remained on the colony with probability *ρ_y_* before leaving the observable colony through dispersal or mortality.

Daily expected abundances were updated recursively. Stage 1 received new births each day, whereas subsequent stages received individuals progressing from the preceding stage while retaining pups already present. Stage 5 accumulated newly weaned pups and lost individuals through departure from the colony according to their annual retention probability. The resulting daily expected abundances formed the latent population process. Conditional on the model parameters, expected daily stage abundances were updated deterministically; stochastic variation between expected and observed stage counts was represented by the Poisson observation model.

Observed stage-specific pup counts were modelled as Poisson random variables with expectation equal to the predicted stage abundance multiplied by the detection probability (*p*),

$$C_{k,s}\sim\mathrm{Poisson}\text{ ⁣}\left( pN_{y_{k},t_{k},s} \right),$$

Where *C_k,s_* is the observed count of developmental stage $s$ during survey *k*, and *y_k_*, and *t_k_* denote the breeding season and model day of that survey. Detection probability *p* was fixed at one for all developmental stages in the primary analysis.

#### Model priors

Prior distributions were informed by published information wherever available and by empirical summaries of the observed data when suitable external information was unavailable (Table S2).

The annual total number of births, *B_y_*, was assigned a truncated normal prior centred on the maximum total number of pups observed during each breeding season with a common SD. The annual peak of the birth season, *μ_y_*, was assigned a truncated normal prior derived from approximate birth dates reconstructed from the observed stage-specific counts. Approximate birth dates were obtained by subtracting representative developmental ages of 2, 8, 12, 17 and 25 days from observations of Stages 1 to 5, respectively (Jenssen et al., 2010), and calculating the weighted mean birth date for each breeding season. The overall prior mean and SD for annual peaks were then calculated from these estimates. The birth-season spread, *σ_b,y_*, was assigned a weakly informative truncated normal prior with mean of 7.5 days, SD of 4.5 days, and bounds of 0.5 and *T*.

Mean durations of developmental Stages 1 to 4, *d_s_*, were informed by published descriptions of grey seal pup development (Jenssen et al., 2010). Published developmental subclasses were combined to match the field-stage classifications used in Jenssen et al. (2010) (Stage 1 = stages 0, 1 and 1+; Stage 2 = stages 2 and 2+; Stage 3 = stages 3 and 3+; Stage 4 = stages 4 and 4+). Stage boundaries were defined as the midpoint between the mean ages of adjacent morphological stages, and prior mean durations were calculated as the interval between successive boundaries.

Annual preweaning survival, *ϕ_y_*, was estimated independently for each breeding season. Identical normal priors were assigned on the logit scale, corresponding to a mean preweaning survival of 0.789 and SD of 0.08 on the probability scale, based on published estimates from Baltic grey seal land-breeding colonies (Jüssi et al., 2008).

Annual mean Stage 5 residence time on the colony, *τ_y_*, was estimated independently for each breeding season using identical truncated normal priors with a mean of 21 days, SD of 6 days, and bounds of 5 to 50 days, informed by published observations of the duration that recently weaned grey seal pups remain ashore before departing to sea (Noren et al., 2008).

Detection probability, *p*, was fixed at one for all developmental stages in the primary analysis, reflecting the high visibility of pups in high-resolution aerial imagery. The influence of this and other modelling assumptions was evaluated through a series of sensitivity analyses.

#### Model fitting

The population model was implemented in R (version 4.5.3) (R Core Team, 2024) using the *nimble* package (NIMBLE Development Team, 2026). Posterior distributions were sampled using three Markov chain Monte Carlo (MCMC) chains, each run for 100,000 iterations. The first 30,000 iterations of each chain were discarded as burn-in, and all remaining iterations were retained, yielding 210,000 posterior samples. Independent initial values were generated for each chain from distributions centred on the specified prior values while respecting parameter constraints. A sensitivity analysis was carried out to test the robustness of posterior estimates to alternative prior specifications, model assumptions, and survey timing (Supplementary materials 1).

Posterior summaries were reported as medians with 95% credible intervals (95% CrIs). Convergence was assessed using the potential scale reduction factor, *R̂*, and effective sample size (ESS) calculated with the *coda* package (Plummer et al., 1999). Model fit was evaluated by comparing posterior predicted stage-specific abundances with the observed survey counts and summarised using mean absolute error (MAE), root mean squared error (RMSE), and the proportion of observations falling within the 95% posterior predictive intervals.

#### Sensitivity analysis

The robustness of posterior parameter estimates to alternative prior distributions and model assumptions was evaluated by refitting the Bayesian state-space population model under a series of alternative prior specifications and detection probabilities. Each sensitivity analysis scenario varied a single model component while holding all remaining priors and model structure unchanged. Scenarios examined the assumed detection probability, the prior mean and uncertainty of developmental stage durations, Stage 5 residence time, preweaning survival, total annual births, birth-season peak, and birth-season spread. Alternative priors for preweaning survival and Stage 5 residence were applied identically and independently to each breeding season. An additional sensitivity analysis evaluated the influence of survey timing by repeating the analysis after excluding all observations collected before the first survey date common to both breeding seasons. Initial abundance was fixed at zero in all sensitivity analyses and was therefore not varied. Sensitivity models were fitted using three MCMC chains of 30,000 iterations, with the first 15,000 iterations discarded as burn-in and the remaining samples thinned by retaining every tenth iteration.

For each scenario, posterior medians and 95% CrIs were extracted for annual preweaning survival and Stage 5 residence time. The influence of each modelling assumption was evaluated by comparing posterior estimates from the alternative scenarios with those obtained under the primary model.

The sensitivity analyses indicated that interannual patterns in posterior estimates of preweaning survival and Stage 5 residence were generally robust to alternative prior distributions and model assumptions (Figures S2 and S3). Preweaning survival was most sensitive to variation in its own prior distribution, whereas changes in the prior distributions of other parameters produced comparatively modest changes. Stage 5 residence estimates were similarly robust to changes in prior distributions, although the 2026 estimate was more sensitive to variation in the Stage 5 residence prior. Across all scenarios, the relative patterns between breeding seasons were unchanged, with similar preweaning survival between years and consistently longer Stage 5 residence in 2026. Restricting the analysis to surveys conducted after the first survey date common to both breeding seasons resulted in modestly higher preweaning survival (0.70 [95% CrI = 0.56 to 0.84] for 2025 and 0.73 [95% CrI = 0.61 to 0.84] for 2026) and slightly shorter Stage 5 residence (10.5 days [95% CrI = 8.4 to 13.7] for 2025 and 28.3 days [95% CrI = 21.4 to 37.0] for 2026), but did not alter these interannual patterns, indicating that the earlier start of the 2026 monitoring programme did not influence the principal conclusions of the analysis.

### Body condition assessment

#### Image processing

A novel object detector employing YOLOv8 was developed to identify grey seals within orthomosaic imagery (Jocher et al., 2026). The detector classified four categories: adults, white-coat pups (Stages 1 to 3), moulting pups (Stage 4), and grey-coat pups (Stage 5). The detector was trained using the Ultralytics Hub platform on 278 individual orthomosaic tiles containing 1314 annotated seal outlines (436 adults, 461 white-coat pups, 111 moulting pups, and 306 grey-coat pups). Images were randomly split between training (70%), validation (20%), and testing datasets (10%). Model performance was evaluated using precision, recall, and F1-score (Infantes et al., 2022; Carroll et al., 2025b). The trained detector was applied to orthomosaics using the Deepness plugin (Aszkowski et al., 2023) within QGIS (version 3.44.11) (Nyall Dawson et al., 2026). The detector was applied with a resolution of 1 cm/pixel, tile overlap of 20%, and required confidence level of 50%. Only Stage 4 detections were retained for body-condition analysis. All detections were visually inspected, with incorrectly classified individuals or individuals poorly represented by the polygon outline (e.g., due to being partially obscured by vegetation) being removed. For each orthomosaic, the numbers of automatic detections, retained detections after quality control, and false-positive detections arising from misclassified pups, adults, non-seal objects, or poor polygon fits were recorded and compared with manual orthomosaic counts.

The resulting Stage 4 polygons were imported into R, where polygon area was calculated using the *st_area()* function from the *sf* package (Pebesma, 2018).

#### Mass estimation

Ground-truth mass measurements were obtained as part of a concurrent field study. Stage 4 pups were captured and weighed in a canvas bag using a hanging balance. True body mass was determined by subtracting the known mass of the bag from the combined mass of the pup and bag. Pups were then marked with a unique letter or number using livestock marker before being released. Targeted drone imagery was collected shortly afterwards, allowing marked individuals to be identified within orthomosaics and their corresponding polygon areas extracted. From images, pups body posture was classified as prone, supine, or lateral recombinant. As only a single pup was identified in supine body posture, this class was not included in subsequent analysis.

The relationship between polygon area and measured body mass was modelled in R. Candidate models were first fitted using linear mixed-effects models implemented in the *lme4* package (Bates et al., 2015). The full model included the fixed effects of polygon area, coat condition, and body posture, together with interactions between polygon area and both coat condition and body posture. Survey date was included as a random intercept to account for variation among drone surveys. A reduced mixed-effects model containing polygon area as the only fixed effect, with survey date retained as a random intercept, was also fitted. The two mixed-effects models were compared using a likelihood-ratio test, and the significance of individual fixed effects in the full model was assessed using Type II analysis of variance implemented in the *car* package (Fox, 2002). As neither coat condition, body posture, their interactions with polygon area, nor the random intercept of survey date significantly improved model fit, the final calibration model was fitted using ordinary least-squares linear regression with polygon area as the sole predictor of body mass. Individual body masses were then predicted from polygon area using the fitted linear model. These estimated masses were subsequently used in all analyses of pup body condition.

The area-to-mass calibration was based on 34 pups of known mass, spanning 14.9 to 58.3 kg and polygon areas of 0.130 to 0.405 m^2^. Polygon area was strongly related to known pup mass (*β* = 151.8 kg/m^2^, standard error = 12.5 kg/m^2^, t-value = 12.13, p-value < 0.001). The mean absolute error was 3.4 kg (Figure S4).

Adding coat stage, body posture, and their interactions with polygon area did not improve model fit relative to the area-only mixed model (χ^2^ = 5.84, df = 7, p-value = 0.558). Coat stage (p-value = 0.139), body posture (p-value = 0.466), area-by-coat interaction (p-value = 0.896), and area-by-posture interaction (p-value = 0.950) were unsupported. Consequently, as the survey-date random effect was negligible, the final calibration used to predict individual pup mass was a simple linear model relating measured body mass to polygon area. Individual mass uncertainty was represented using 95% prediction intervals.. Estimated masses for the 137 Stage 4 pups ranged from 21.7 to 51.4 kg, with an overall mean of 37.5 with a SD of 6.6 kg. Across surveys, mean estimated mass was 37.1 with a SD of 6.4 kg in 2025 and 38.0 with a SD of 6.9 kg in 2026.

#### Cumulative density exposure

Pup density was estimated as number of pups present per 100 m² of breeding area. Following Jüssi et al. (2008), the breeding area was assumed to be constant and comprise 75% of the 8,000 m² area of Innarahu (6,000 m²). To estimate the breeding environment experienced by pups during development, date of birth was approximated from survey date using the posterior median estimates of developmental stage duration obtained from the fitted state-space population model. Moulting pups were assumed to have been surveyed at the midpoint of Stage 4 (age = *d_1_* + *d_2_* + *d_3_* + *d_4_/2*).

Because daily density was expressed as pups per 100 m² and summed across daily time steps, cumulative density exposure was expressed as pup-days per 100 m². Cumulative density exposure was calculated from posterior estimates of daily pup abundance, producing posterior distributions for each survey occasion. Posterior medians were used as fixed survey-level predictors in the body-mass analysis, while posterior uncertainty was retained when calculating credible intervals for the density exposure metrics. These metrics represent population-level indices of the colony environment experienced during development rather than exact individual exposure histories.

### Supplementary tables

**Table S1. Classification criteria used to assign grey seal pups to developmental stages during aerial-image analysis.** Adapted from Jenssen t al. (2010).

| Stage | Short description | Diagnostic characteristics |
| --- | --- | --- |
| 1 | Thin white coat | Neck well defined and loose skin folds present around the body. A fleshy pink umbilicus may be visible if the ventral surface is exposed. The lanugo is intact, although a limited area of the muzzle may be exposed, and may be white or stained yellow or red by birth fluids. |
| 2 | Intermediate white coat | Body contour smoother than Stage 1 with no loose skin folds, although the neck remains visible. The umbilicus is absent if the ventral surface is exposed. The lanugo is intact, although a limited area of the muzzle may be exposed. |
| 3 | Fat white coat | Body contour rounded and barrel-shaped with no visible neck. The lanugo is intact, although a limited area of the muzzle may be exposed. |
| 4 | Moulting | Body condition is variable. The lanugo has begun to moult beyond the muzzle, revealing the underlying silver, grey or black pelage. At least 5% of the visible body surface remains covered by lanugo. |
| 5 | Grey coat | Less than 5% of the visible body surface remains covered by lanugo. |

**Table S2. Prior distributions used in the Bayesian state-space population model.** Prior distributions are given for all estimated model parameters, together with the parameter values used in the primary analysis and the source or rationale used to specify each prior. Parameters derived deterministically within the model (e.g. daily stage-transition probabilities and stage-specific survival probabilities) are not included because they were not assigned independent prior distributions.

| Parameter (unit) | Notation | Distribution | Prior values | Source |
| --- | --- | --- | --- | --- |
| Annual total births (pups) | *B_y_* | Truncated Normal | 2025: Mean = 219; 2026: Mean = 159, SD = 42.4 Lower bound = 0 | Empirical summary of observed counts |
| Birth peak (days) | *μ_y_* | Truncated Normal | Mean = 15.8 SD = 1.90 Bounds = 1 to *T* | Approximate birth dates reconstructed from observed stage counts |
| Birth spread (days) | *𝜎_y_* | Truncated Normal | Mean = 7.5 SD = 4.5 Bounds = 0.5 to *T* | Weakly informative prior |
| Stage 1 duration (days) | *d_1_* | Truncated Normal | Mean = 6.5 SD = 1.5 Bounds = 3 to 10 | (Jenssen et al., 2010) |
| Stage 2 duration (days) | *d_2_* | Truncated Normal | Mean = 4.5 SD = 1.5 Bounds = 2 to 8 | (Jenssen et al., 2010) |
| Stage 3 duration (days) | *d_3_* | Truncated Normal | Mean = 5.5 SD = 2.0 Bounds = 2 to 10 | (Jenssen et al., 2010) |
| Stage 4 duration (days) | *d_4_* | Truncated Normal | Mean = 7.0 SD = 2.5 Bounds = 3 to 14 | (Jenssen et al., 2010) |
| Annual preweaning survival (probability) | *ϕ_y_* | Normal | On logit scale, centred on probability = 0.789  SD = 0.08 | (Jüssi et al., 2008) |
| Annual Stage 5 residence (days) | *τ_y_* | Truncated Normal | Mean = 21 SD = 6 Bounds = 5 to 50 | (Noren et al., 2008) |
| Detection probability | *p* | Fixed | 1.00 | Assumed in primary analysis |

**Table S3. Sensitivity-analysis scenarios evaluated in the Bayesian state-space population model.** Each sensitivity analysis modified a single model assumption while holding all remaining priors, model structure, and initial conditions unchanged. The full model was refitted under each alternative scenario, and posterior estimates were compared with those obtained under the primary analysis.

| Sensitivity component | Baseline value | Alternative scenarios |
| --- | --- | --- |
| Detection probability | 1.00 | 0.80, 0.90, 0.95, 0.99 |
| Stage-duration prior means | 6.5, 4.5, 5.5, 7.0 days | 0.85 × baseline; 1.15 × baseline |
| Stage-duration prior SDs | 1.5, 1.5, 2.0, 2.5 days | 1.5 × baseline; 2.0 × baseline |
| Stage 5 residence prior mean | 21 days | 14, 18, 24, 28 days |
| Stage 5 residence prior SD | 6 days | 3, 9, 12 days |
| Preweaning survival prior mean | 0.789 | 0.65, 0.70, 0.75, 0.83, 0.88 |
| Preweaning survival prior SD | 0.08 | 0.04, 0.12, 0.16 |
| Total births prior mean | 2025: 219; 2026: 159 pups | 0.85 × baseline; 1.15 × baseline |
| Total births prior SD | 2025: 42.1; 2026: 42.4 pups | 0.75 × baseline; 1.50 × baseline |
| Birth-peak prior mean | 15.8 model day | Baseline ±2 and ±4 model days |
| Birth-peak prior SD | 1.90 days | 0.5 × baseline; 1.5 × baseline |
| Birth-spread prior mean | 7.5 days | 4.5, 6.0, 9.0, 10.5 days |
| Birth-spread prior SD | 4.5 days | 2.5, 6.5 days |
| Survey timing | All available surveys | Exclude surveys conducted before the first survey date common to both years |

**Table S4. Performance of automated image segmentation for identifying Stage 4 (moulting) grey seal pups.** *Manual* is the number of Stage 4 pups identified by manual counts. *Auto*. is the total number of polygons returned by machine learning classification. *Retained* is the number of Stage 4 pups retained for body-mass analysis after manual quality control. *False pos. (Miss.)* are seals which were incorrectly classified as Stage-4 by during automated classification. *False pos. (True)* are detections of non-seal objects. *Poor fit* are polygons correctly identified as Stage 4 pups but removed due to poor representation of the seal outline.

| Year | Date | Manual | Auto. | Retained | | False pos. (Miss.) | False pos. (True) | Poor fit |
| --- | --- | --- | --- | --- | --- | --- | --- | --- |
| 2025 | 25 Feb | 17 | 29 | 13 | Adult: 2  Pup: 14 | | 0 | 0 |
| 2025 | 5 Mar | 45 | 55 | 37 | Adult: 0  Pup: 15 | | 1 | 2 |
| 2025 | 13 Mar | 60 | 34 | 20 | Adult: 0  Pup: 10 | | 0 | 4 |
| 2026 | 25 Feb | 10 | 7 | 7 | Adult: 0  Pup: 0 | | 0 | 0 |
| 2026 | 5 Mar | 34 | 30 | 28 | Adult: 1  Pup: 1 | | 0 | 0 |
| 2026 | 13 Mar | 39 | 34 | 32 | Adult: 0  Pup: 2 | | 0 | 0 |
| total | — | 205 | 189 | 137 | Adult: 3  Pup: 42 | | 1 | 6 |

### Supplementary figures

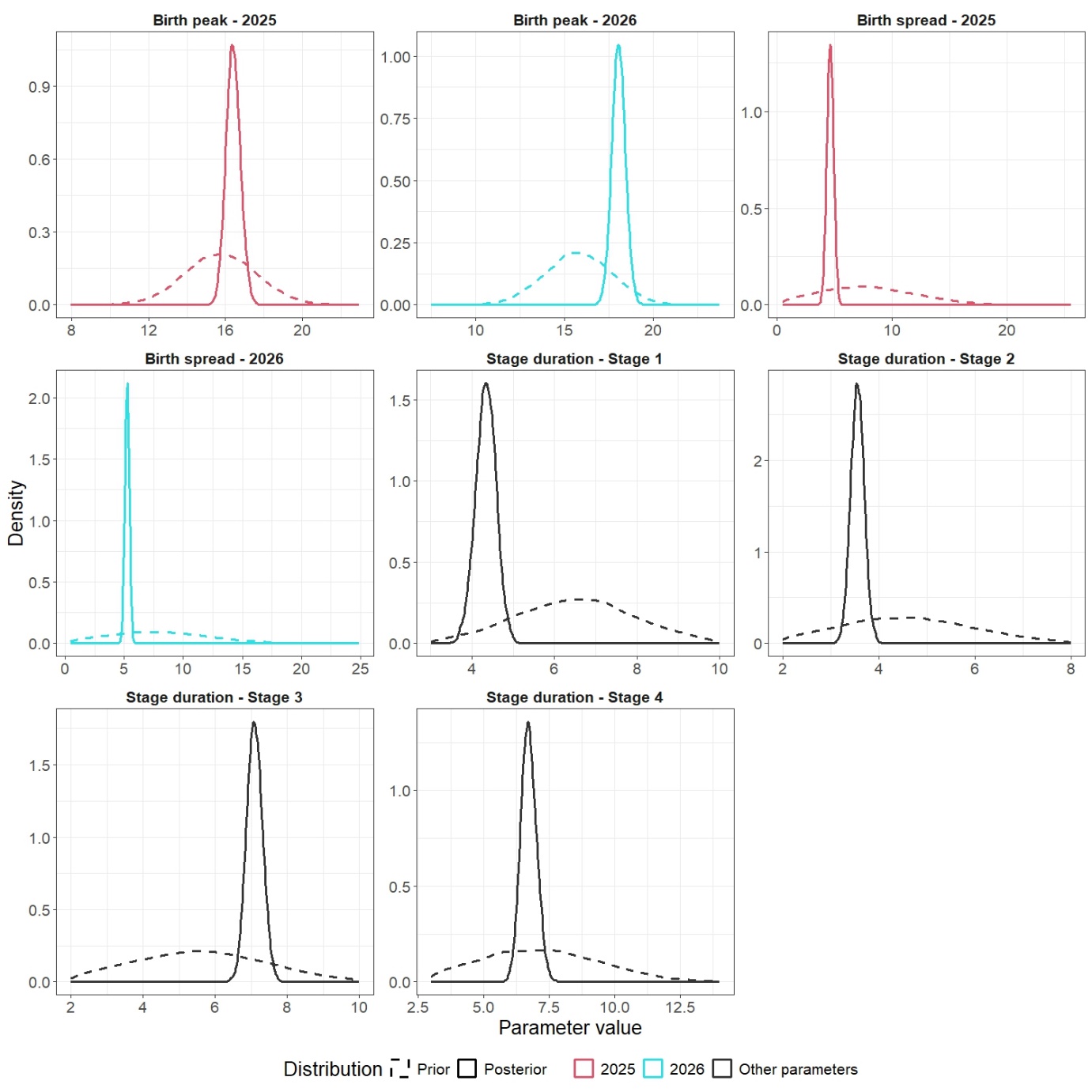

**Figure S1. Prior and posterior distributions of the remaining parameters estimated by the Bayesian state-space population model.** Prior (dashed lines) and posterior (solid lines) distributions are shown for annual birth peak, annual birth spread, and developmental stage durations. Year-specific parameters are coloured by breeding season (2025, red; 2026, blue), whereas parameters common to both years are shown in black.

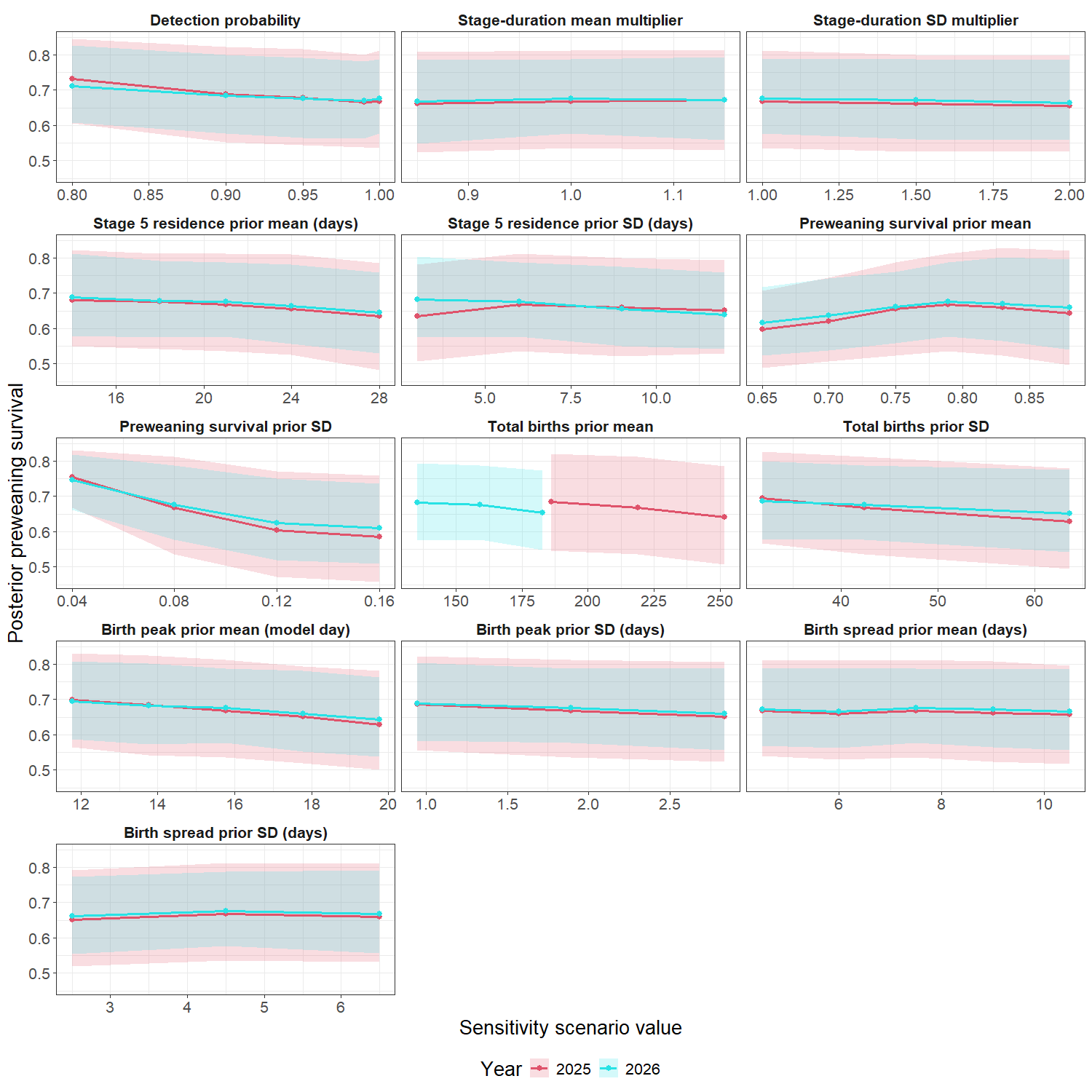

**Figure S2. Sensitivity of posterior annual preweaning survival estimates to alternative prior values and model assumptions.** Each panel shows posterior median preweaning survival (solid lines) and 95% CrIs (shaded ribbons) for the 2025 and 2026 breeding seasons after refitting the Bayesian state-space population model under alternative sensitivity scenarios. In each analysis, a single model prior parameter was varied while all remaining priors, model structure and initial conditions were held constant.

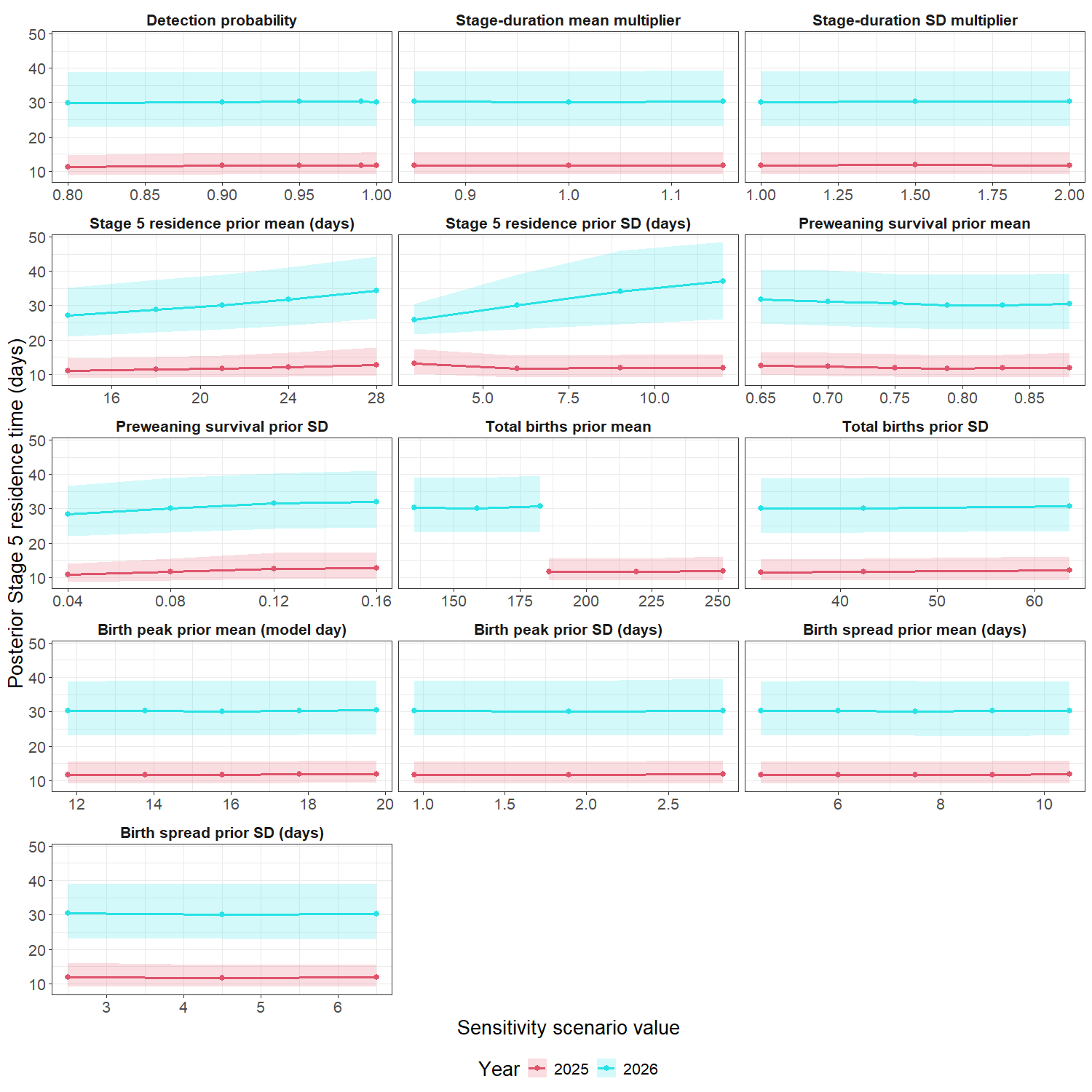

**Figure S3. Sensitivity of posterior annual Stage 5 residence estimates to alternative prior values and model assumptions.** Each panel shows posterior median Stage 5 residence time (solid lines) and 95% CrIs (shaded ribbons) for the 2025 and 2026 breeding seasons after refitting the Bayesian state-space population model under alternative sensitivity scenarios. In each analysis, a single model prior parameter was varied while all remaining priors, model structure and initial conditions were held constant.

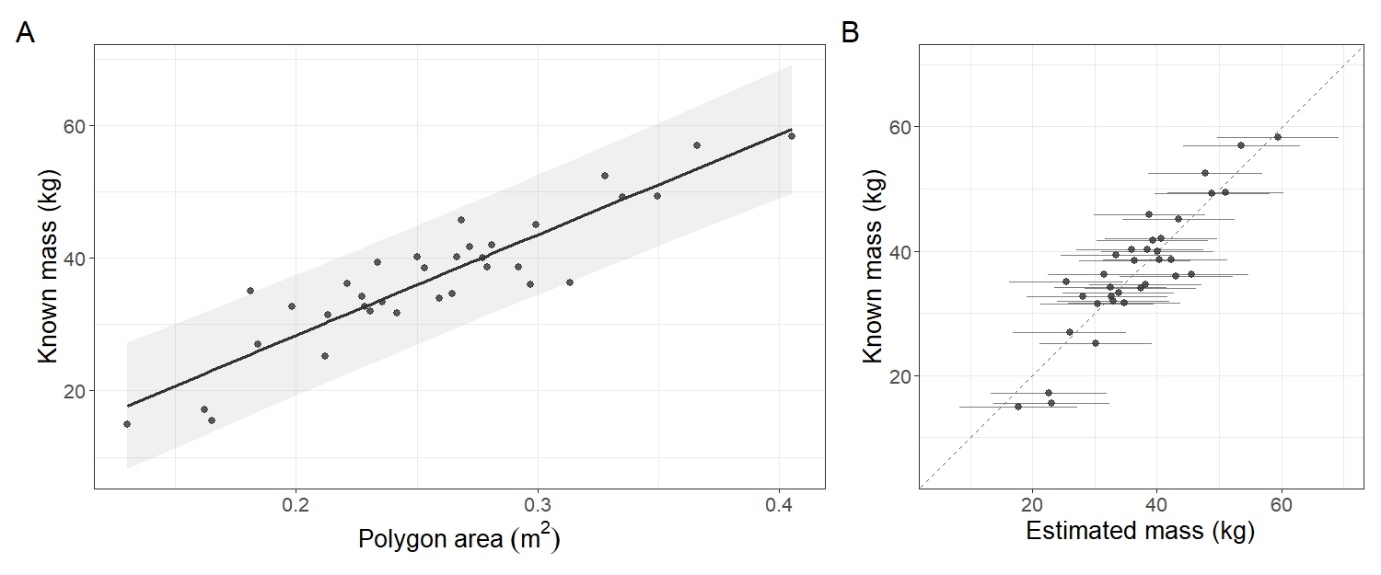

**Figure S4. Calibration of the relationship between polygon area and grey seal pup mass.** (**A**) Linear relationship between pup polygon area and known pup mass used to estimate body mass from drone imagery. Shaded ribbon shows the 95% prediction interval. (**B**) Predicted versus known pup mass for the calibration dataset. Horizontal error bars represent 95% prediction intervals, and the dashed line indicates the one-to-one relationship.

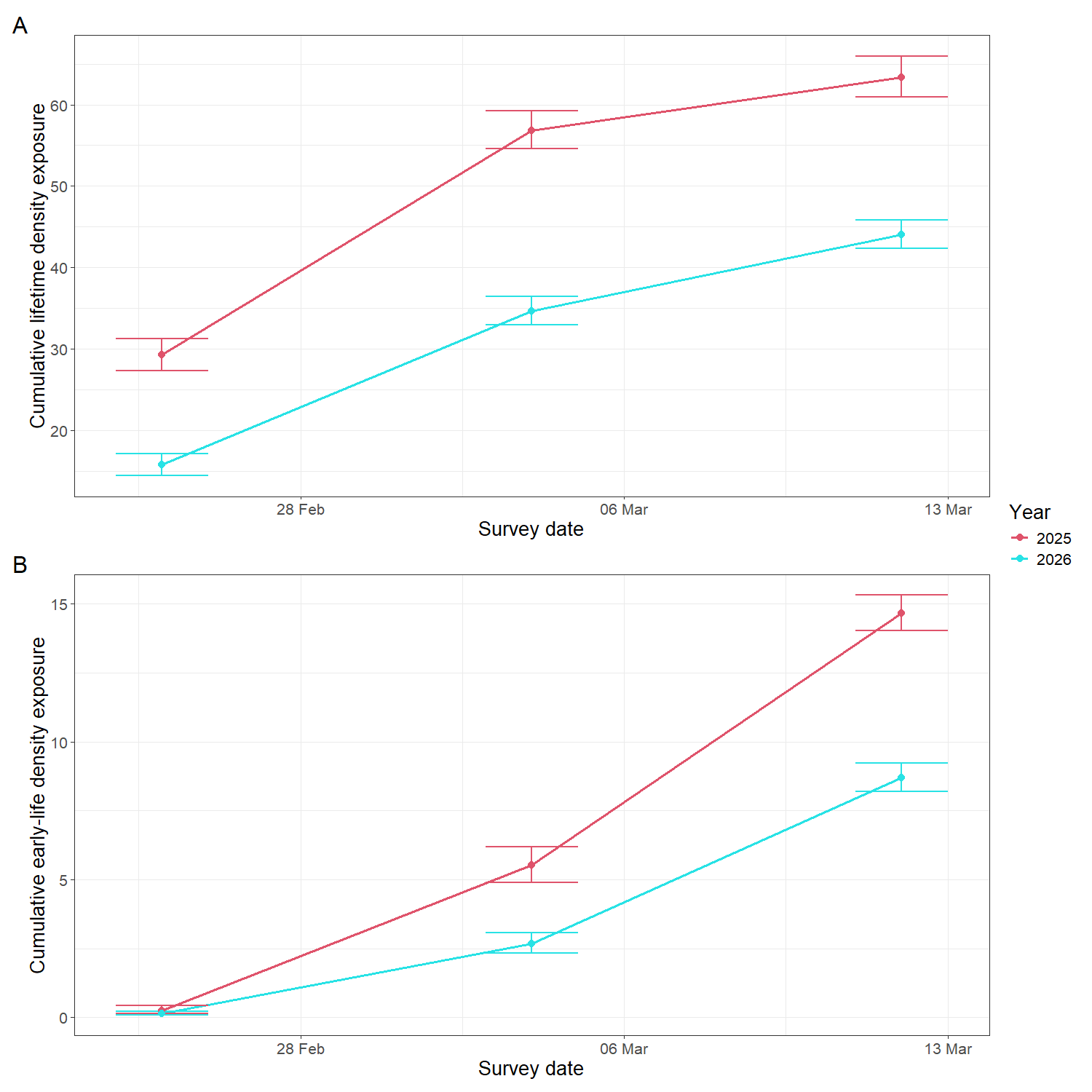

**Figure S5. Cumulative density exposure metrics assigned to moulting grey seal pups from the Bayesian state-space population model.** (**A**) Cumulative lifetime density exposure. (**B**) Cumulative early-life density exposure. Points show the median cumulative density exposure calculated across posterior draws of daily pup abundance for each survey occasion, and vertical error bars show 95% CrIs. Uncertainty reflects posterior uncertainty in daily pup abundance propagated through the calculation and temporal accumulation of pup density.
